# Single molecule studies reveal that p53 tetramers dynamically bind response elements containing one or two half sites

**DOI:** 10.1101/2019.12.19.883140

**Authors:** Elina Ly, Jennifer F. Kugel, James A. Goodrich

## Abstract

The tumor suppressor protein p53 is at the nexus of cell fate decisions, including apoptosis, senescence, and cell cycle arrest. p53 is a tetrameric transcription factor that binds to DNA response elements to regulate transcription of its target genes, a process activated by cellular stress. p53 response elements consist of two decameric half-sites, and most data suggest one p53 dimer in the tetramer binds to each half-site. Despite a broad literature describing p53 binding to DNA, unanswered questions remain, due in part to the need for more quantitative and structural studies with the full length protein. Here we describe a single molecule fluorescence system to visualize full length p53 tetramers binding to DNA in real time. The data reveal a dynamic interaction with many p53 binding and dissociation events occurring on single DNA molecules over minutes. We found that p53 tetramers bound to response elements containing only a single half site. The kinetic stability of tetramer/DNA complexes depended on the number of half sites and the helical phasing between them, with the most stable complexes forming on DNA containing two adjacent half sites. The forward rate of binding was not strongly impacted when one half site was mutated. These studies provide real time kinetic measurements of full length p53 tetramers binding to single molecules of DNA, and reveal new insight into the mechanisms by which this nucleoprotein complex forms.

## Introduction

Transcriptional activators control when and in what cell type specific genes are expressed, a regulatory process that underpins all aspects of cell biology. p53 is a critical transcriptional activator that controls cell fate decisions including apoptosis, senescence, and cell cycle arrest in response to stresses such as DNA damage (1, 2). Because p53-dependent transcriptional pathways are activated in response to genotoxic stress, p53 is often referred to as the guardian of the genome (3, 4). Indeed, p53 mutations – often impairing its ability to bind to DNA response elements (REs) – result in carcinogenic phenotypes across a broad spectrum of cell types (5), and p53 is one of the most commonly mutated genes in human cancers (6, 7). p53 is also important for regulating cellular metabolism in response to low levels of constitutive stress (2). In all cases, the ability to activate transcription is key to how p53 functions, underscoring the importance of unraveling the molecular mechanisms by which it activates transcription.

To activate transcription, p53 must recognize and bind to p53 REs in DNA. The p53 RE is composed of two 10 bp half site sequences with 0 to 13 base pairs of spacing in between (8, 9). Each half site is defined by the palindromic consensus sequence RRRCWWGYYY, where R represents A or G, W represents A or T, and Y represents C or T (8, 9). A p53 RE can be located proximal to the core promoter of a gene it regulates, or many kilobases away within enhancer regions (10). Genome-wide data sets interrogating p53 occupancy reveal hundreds to thousands of binding sites, depending on the cell type and conditions (11–13). Our understanding of how p53 binds its RE has been informed from numerous important biochemical and structural studies that describe unifying principles (14); however inconsistencies and questions remain. A contributing factor is the need for more structural and quantitative biochemical studies using the full length p53 protein, particularly in light of evidence showing that regions outside the DNA binding domain can impact the interaction of p53 with DNA (15–20).

The domain structure of p53 consists of an acidic N-terminal region, a core DNA binding domain (DBD), an oligomerization domain (OD), and an unstructured basic C-terminal domain (CTD) (14). The N-terminal region contains two unstructured transcriptional activation domains that interact with co-regulatory complexes to mediate transcriptional activation (21). Recent work revealed that intramolecular interactions between the N-terminal region and the DBD can impact the affinity and specificity with which p53 binds DNA (19, 20). The DBD is largely responsible for recognizing and making contacts with p53 REs. It contains the highest frequency of cancer associated mutations, highlighting the importance of DNA binding to the function of p53 (22). The OD facilitates the formation of the p53 tetramer, commonly thought of as a dimer of dimers. Tetrameric p53 is considered the fully active DNA-bound form (23–25); however, some data suggest that dimeric p53 can trigger specific transcriptional programs in cells (26, 27). The role of the basic unstructured CTD is more enigmatic, and the majority of biochemical studies of p53 DNA binding use versions of the protein lacking the CTD. Initially the CTD was thought to negatively regulate DNA binding, but more recent work has shown the CTD can stabilize p53 binding to REs as they diverge from consensus (15, 16), and also facilitate DNA sliding as p53 searches for its specific binding sites (17, 18).

Crystal structures of the isolated DBD bound to a single DNA half site revealed two DBD monomers bound per half site, with two such dimer/DNA complexes packing into tetramers, leading to the model that p53 binds DNA as a dimer of dimers (28). The structure of the p53 DBD bound to a complete RE (i.e. two half sites) also revealed tetramers binding DNA (one dimer per half site) with extensive protein-protein contacts occurring at the interface between the two dimers(29). To date high resolution structures of full length p53 bound to DNA have not been solved. Lower resolution EM studies with full length p53 suggest a p53 tetramer can bind a single half site, and moreover, that an octamer can assemble on the DNA if p53 tetramers bind both half sites present in an RE (30–32).

To provide insight into the fundamental mechanisms by which p53 binds to DNA, we developed a single molecule assay to measure binding and dissociation of full length human p53 on DNA molecules in real time. We investigated binding to the p53 RE from the GADD45 promoter, as well as mutations of this RE that eliminated one of the consensus half sites or changed the spacing between half sites. We found that p53 tetramers dynamically bind to and dissociate from DNA, with many of these events occurring over minutes. Our data show that p53 tetramers can bind to DNAs containing only a single half site, however, the kinetic stability of this complex is reduced compared to tetramers binding to a full p53 RE. Moreover, if the spacing between half sites puts them on opposite sides of the DNA helix, tetrameric p53 binds as if there were only a single half site present. Our studies report real time kinetic measurements of p53 tetramers binding to DNA and suggest new mechanisms by which this interaction occurs.

## Results

### Full-length p53 labeled with AF647 tightly binds DNA containing one or two half-sites

With the goal of visualizing p53 binding to DNA using single molecule microscopy, we generated full length human p53 with a C-terminal SNAP tag (p53-SNAP) for fluorescent labeling (33). p53-SNAP was expressed in insect cells then purified via a His tag present on the N-terminus of the protein. We compared p53-SNAP and full length wild type p53 binding to the p53 RE from the GADD45 promoter using electrophoretic mobility shift assays (EMSAs) (Figure 1A). The addition of the SNAP tag did not appreciably impact the DNA binding activity of p53. Next, the p53-SNAP protein was fluorescently labeled with an AlexaFluor647 SNAP dye substrate (AF647-p53). We determined there was near complete coupling of the AF647 dye to p53-SNAP using LC MS/MS. We also evaluated the extent of labeling using fluorescence and absorbance, which each indicated ∼70% labeling efficiency. Therefore, we concluded that ∼30% of the conjugated AF647 dyes were not photoactive, which is consistent with our previous observations (34).

**Figure 1.**
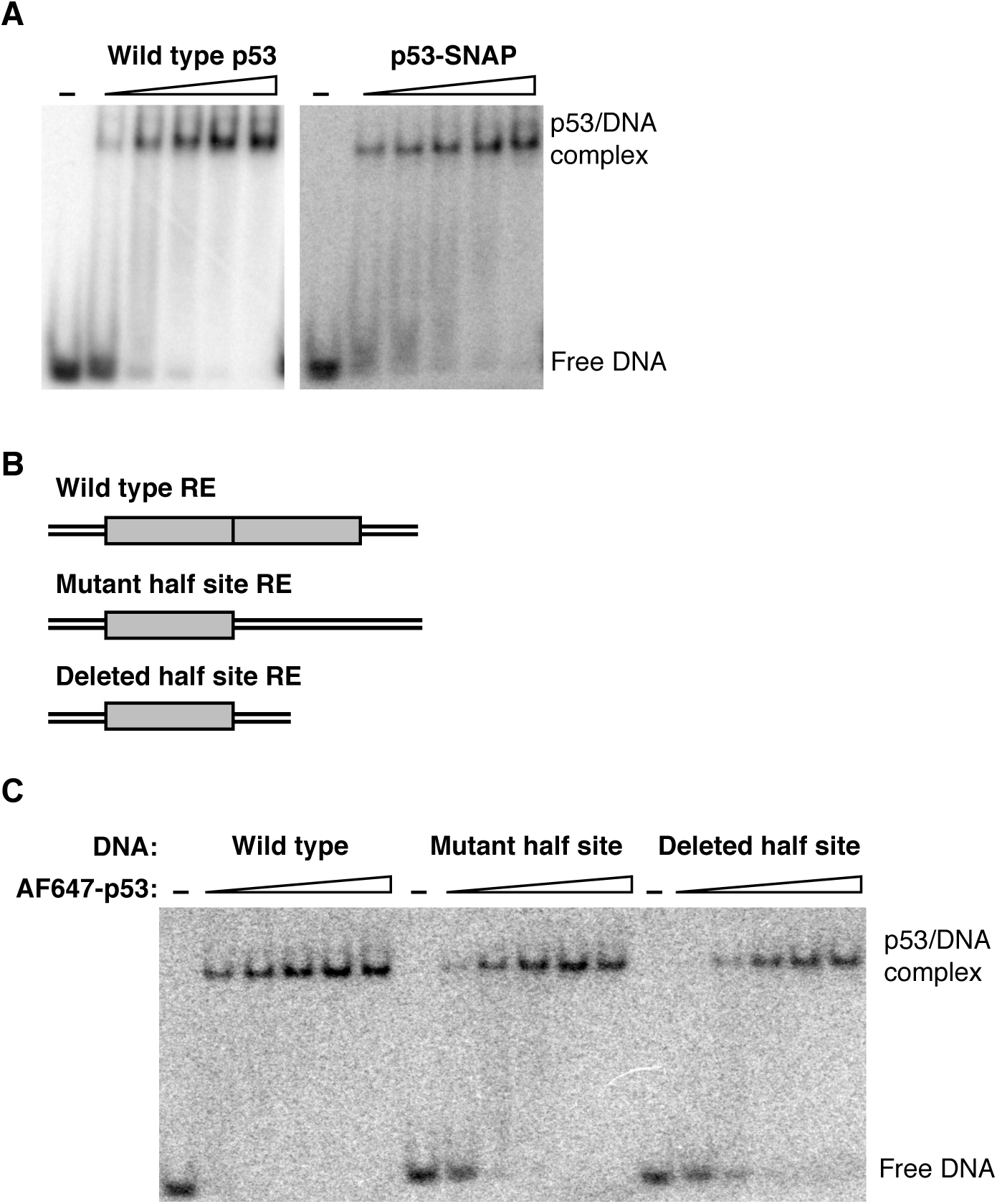
Full length p53 labeled with AF647 via a SNAP tag binds DNA with sub-nM affinity in EMSAs. **A**. The addition of a C terminal SNAP tag to full length p53 does not appreciably alter DNA binding activity. The wild type p53 and p53-SNAP titrations were as follows (tetramer concentrations): 0.025, 0.075, 0.25, 0.75, and 2.5 nM. In all experiments p53 concentrations are expressed as maximum tetrameric concentration, given the monomer concentrations added to the assays. **B**. Schematics showing the half site composition of the p53 REs tested. The wild type 32 bp DNA is from the GADD45 promoter and consists of two 10 bp half sites (gray boxes) with no spacing between them. The mutant half site DNA was the same length (32 bp) and had one half site scrambled, whereas the deleted half site DNA had one half site removed and was therefore shorter (22 bp) and. **C**. p53 binds DNA containing one or two half sites with high affinity and the same oligomeric state. The AF647-p53 titration was as follows (tetramer concentration): 0.075, 0.25, 0.75, 2.5, and 7.5 nM.

We tested the ability of AF647-p53 to bind the three DNA constructs diagrammed in Figure 1B using EMSAs. The wild type DNA consisted of the RE from the human GADD45 promoter, containing two 10 bp half-sites (gray boxes) with no spacing between. The mutant half site construct had one half site sequence randomized and the other remained intact. The deleted half site construct was missing one half site, and therefore was 10 bp shorter in length compared to the other two constructs. These DNAs were ^32^P-labeled and incubated with increasing concentrations of AF647-p53, then free versus bound DNA was resolved in a native gel (Figure 1C). Full length p53 bound the wild type DNA with sub-nM affinity, assuming it bound as a tetramer, which is consistent with most models. When one half site was mutated or removed, the apparent affinity decreased slightly. Even with these decreases, however, p53/DNA complexes still formed on the single half-site DNAs with apparent sub-nM affinity. The most striking observation from these data is that the bound DNA complexes on the three different DNAs migrated at identical positions in the gel. This indicates that p53 binds the wild type DNA and the single half site DNAs with the same oligomeric state, presumably a tetramer.

### A single molecule fluorescence microscopy assay to visualize DNA binding by p53 tetramers

We next used AF647-p53 to establish a TIRF (total internal reflection fluorescence) microscopy system to quantify full length p53 binding to DNA in real time at the single molecule level. This would allow us to directly determine whether p53 binds single half site DNAs as a tetramer, and also measure how the kinetics of p53 binding is impacted by the number of half sites. As illustrated in Figure 2A, biotinylated DNA constructs labeled with an AF647 fluorophore were immobilized on the surface of slides functionalized with streptavidin. To mark the positions of AF647-labeled DNA on the surface, images of red emission were obtained with a TIRF microscope (DNA movie). Subsequently, AF647-p53 was flowed into the slide chambers and incubated with the DNA for 10 minutes. A red emission movie was recorded over the same regions of the slide (DNA+p53 movie). Spots of AF647 emission from the DNA movie were co-localized with spots of AF647 emission from the DNA+p53 movie. For each spot pair the fluorescence emission over time was followed to identify binding and dissociation events. Importantly, the AF647 on the DNA molecule provided a baseline intensity for a single red dye at that position on the surface. This baseline emission intensity was used to “count” the number of dye molecules associated with each AF647-p53 binding event, thereby revealing its oligomeric state.

**Figure 2.**
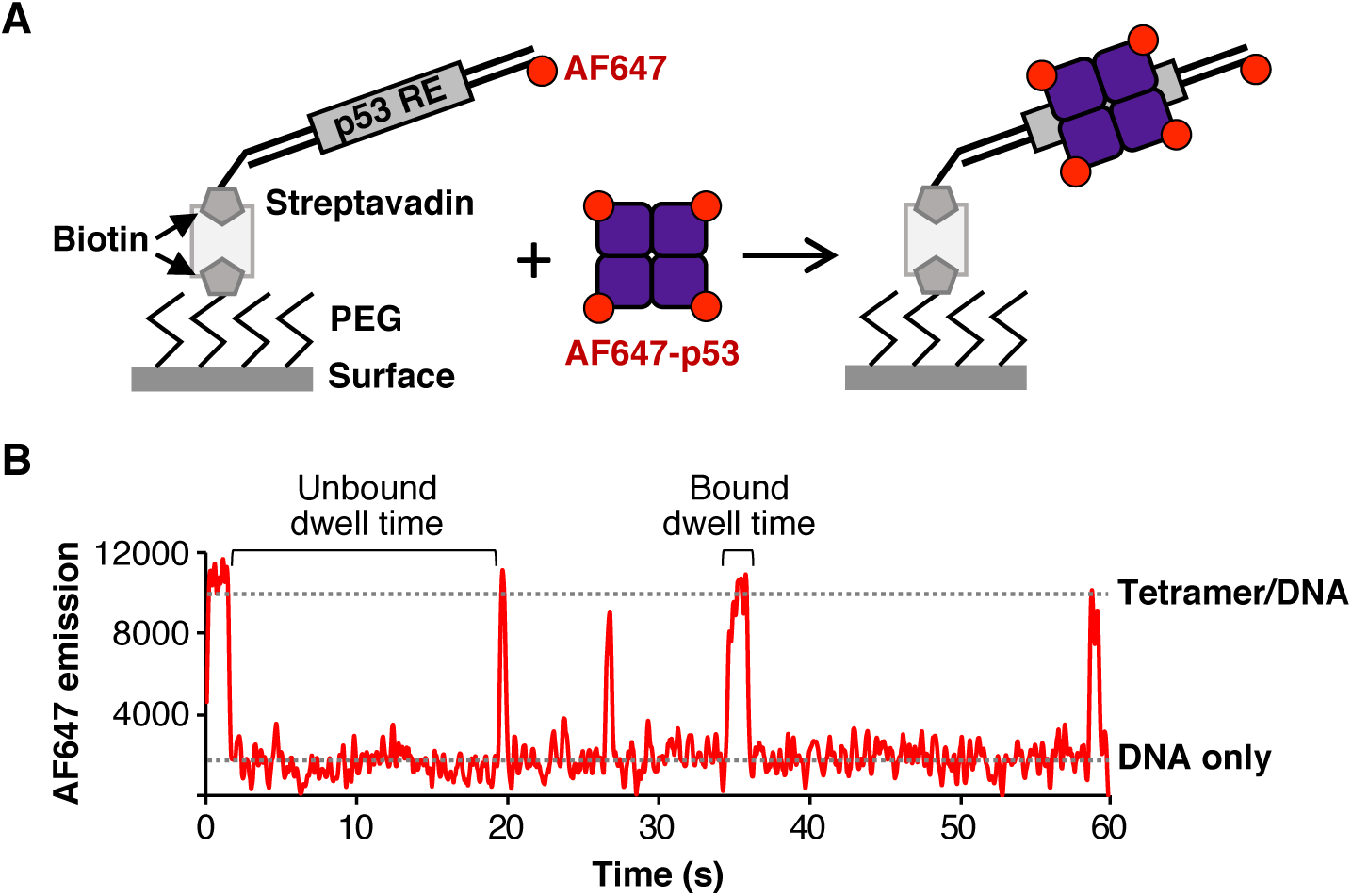
Single molecule fluorescence can be used to detect dynamic binding of tetrameric p53 to immobilized DNA in real time. **A**. Illustration of the single molecule assay used to quantify tetrameric p53 binding to DNA. Red emission movies of biotinylated AF647 labeled DNA immobilized on the streptavidin derivatized surface were collected. AF647-p53 was flowed in and another red emission movie was collected over the same region (p53+DNA). Co-localization of molecules in the DNA only data and the p53+DNA data identified spot pairs from which emission intensity traces over time were extracted. These were used to calculate the base intensity value for a single dye on the DNA, and subsequently calculate the number of dyes (i.e. oligomeric state) of the p53 bound to DNA. **B**. Representative AF647 emission data from a single DNA molecule with five tetrameric p53 binding events, shown by the sharp increases in red intensity. Representative regions used to obtain unbound and bound dwell times are indicated.

Figure 2B shows representative data for a single DNA molecule on which five p53 binding events occurred over 1 min, reflected by the five spikes in red emission intensity. The change in fluorescence emission during each of the binding and release events was ∼4-fold greater than the emission from the single red dye on the DNA, showing that each binding and dissociation event involved tetrameric p53. In principle, this approach could allow us to distinguish between dimer versus tetramer binding. However, since only ∼70% of the AF647-p53 molecules contained photoactive dyes, a non-negligible fraction of dimer binding events could be due to partially labeled tetramers (i.e. tetramers containing only two photoactive fluorophores). Therefore, to be confident in our interpretations, we focused our studies only on tetrameric binding events that showed three or four active AF467 dyes binding and dissociating.

### p53 tetramers bind to DNAs containing either one or two half sites

We used our single molecule assay to determine whether p53 tetramers bind to DNAs containing one half site, as suggested by the EMSAs in Figure 1C. Single molecule experiments were performed using AF647-p53 and the three DNAs depicted in Figure 1B, but labeled with AF647 and immobilized on slide surfaces. We also tested a randomized DNA in which the entire p53 RE was scrambled to no longer resemble the consensus sequence. In all single molecule experiments we used 1 nM monomeric p53, which corresponds to a maximum tetrameric concentration of 0.25 nM. For each DNA we counted the number of dynamic tetrameric binding events over 1000 frames, taking data every 60 ms.

As shown in Figure 3A, tetrameric AF647-p53 binding events were observed on the three DNAs with either one or two half sites. The average number of binding events observed on the three DNAs were not statistically different from one another (p values for all pairwise comparisons were > 0.05). The binding events were specific for p53 half sites (be it one or two) because virtually no binding was observed on the random DNA. We also tested the random DNA for p53 binding using EMSAs. As shown in Figure 3B, complexes were observed on the random DNA at low nM concentrations of AF647-p53 tetramers, which is consistent with prior ensemble experiments showing that p53 has nM affinity for random DNA (23, 35). It is likely the caging effect in the native gel trapped kinetically unstable p53/DNA complexes that were not detectable in the single molecule system due to rapid dissociation.

**Figure 3.**
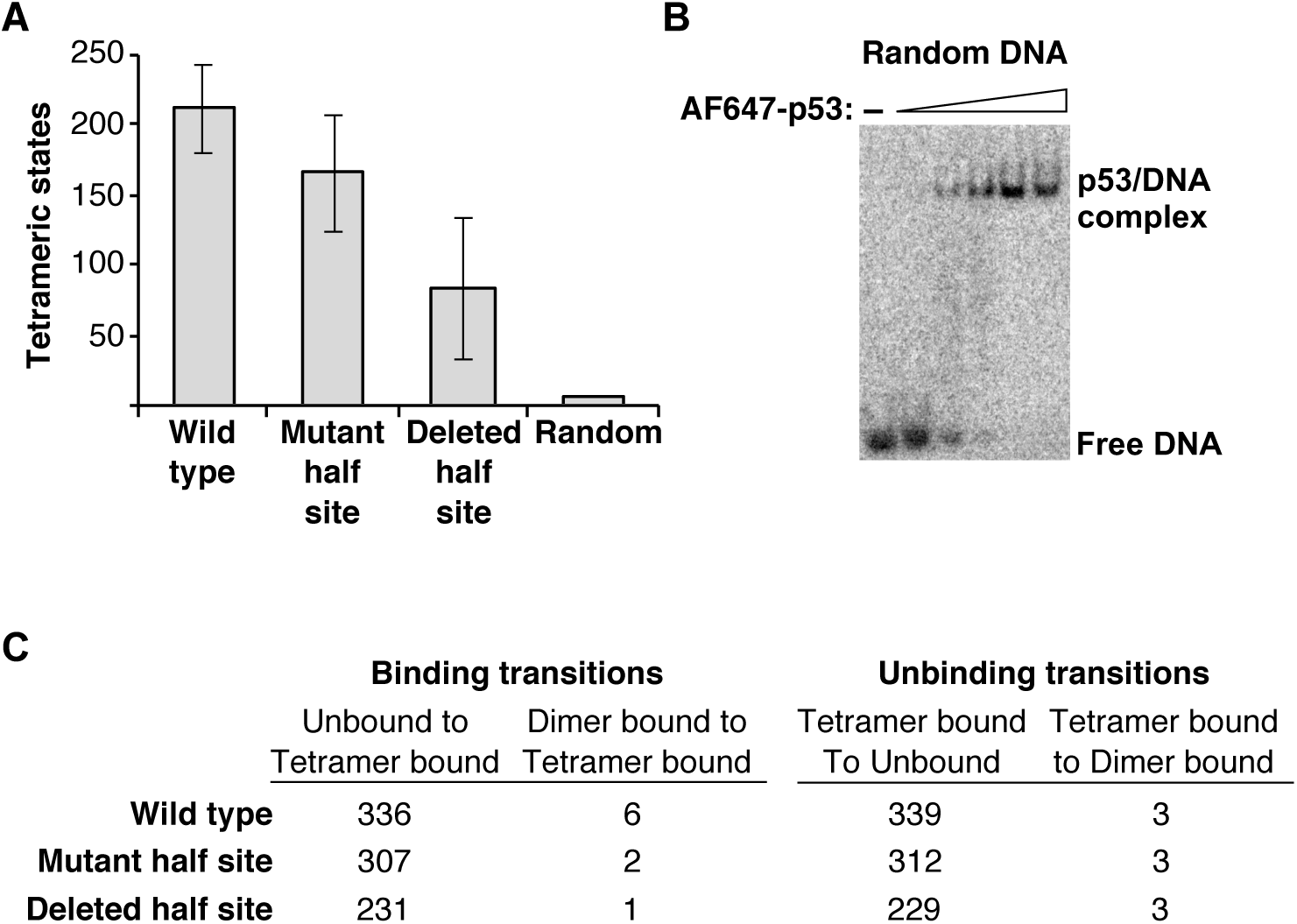
p53 specifically binds as a tetramer to DNA containing one or two half sites. **A**. AF647-p53 binds as a tetramer to wild type DNA and DNAs with a mutated or a deleted half site. The bars represent the average number of dynamic tetrameric states observed from two experiments for the wild type and mutant half site DNAs, and three experiments for the deleted half site DNA. Data with the random DNA were obtained from one experiment over four slide regions. Error bars are the range of the measurements for the wild type and mutant half site DNAs and the standard deviation of the measurements obtained with the deleted half site DNA. **B**. p53 binds to DNA lacking a p53 binding site in EMSAs. The titration of AF647-p53 was as follows (tetramer concentrations): 0.075, 0.25, 0.75, 2.5, and 7.5 nM. **C**. Tetrameric p53 does not go through a dimer/DNA intermediate en route to binding or unbinding. Shown for each DNA are the number of binding and unbinding transitions across two experiments for a total of 8 slide regions.

We also assessed the pathways by which p53 tetramer/DNA complexes formed and dissociated on each DNA molecule. In other words, we evaluated whether tetramer/DNA complexes formed or dissociated via a pathway involving a dimer/DNA intermediate. As shown in Figure 3C, we counted the number of tetrameric binding transitions that showed unbound DNA going to a tetramer-bound state versus the number of dimer-bound states going to tetramer. The opposite transitions were counted for unbinding events. For all three DNA constructs, the vast majority of binding and dissociation events involved the simultaneous addition or loss of ∼4 dyes. Fewer than 2% of the binding and dissociation events involved the gain or loss of two dyes molecules en route to the final bound or unbound state. Hence, we conclude that p53 binding events, regardless of one or two half sites, involve tetramers associating with and dissociating from the DNA molecules.

### p53/DNA complexes are more stable when both half sites are present on the DNA

Given the observation that p53 tetramers bound all three DNAs containing at least one half site, we asked whether there were differences in the kinetics of binding or dissociation. We first measured the observed rate constant for association (k_on(obs)_) and the rate constant for dissociation (k_off_) on the wild type DNA. These measurements were made at a single concentration of protein, hence k_on(obs)_ is presented as an observed first order rate constant. To obtain k_on(obs)_ we measured unbound dwell times (i.e. the time between two p53 tetramer binding events on a single DNA molecule, see Figure 2B). To obtain k_off_ we measured bound dwell times (i.e. the time each tetrameric p53 molecule remained bound to a DNA molecule). The unbound and bound dwell times were histogrammed separately and the data were fit to a single exponential (Figure 4), yielding an average k_on(obs)_ of 0.013 ± 0.002 s^-1^ and an average k_off_ of 0.30 s^-1^ ± 0.06 on the wild type DNA (Table 1). Hence, tetrameric p53/wild type DNA complexes form with a half-time of 52 sec and dissociate with a half-time of 2.3 sec. From these data we calculated a K_D_ for binding wild type DNA of ≤ 5.8 nM (Table 1); our calculations were made assuming all p53 in the reaction is tetrameric, hence the K_D_ values are presented as upper limits.

**Table 1.** A summary of the on and off rate constants, half-times, and apparent K_D_ values for p53 tetramers binding to different DNAs under equilibrium conditions. nd is not determined.

| **DNA** | **$k_{on(obs)}$ (s<sup>-1</sup>)<sup>a</sup>** | **Half-time**<br>**binding (s)<sup>b</sup>** | **$k_{off}$ (s<sup>-1</sup>)<sup>a</sup>** | **Half-time**<br>**unbinding (s)<sup>b</sup>** | **$appK_D$**<br>**(nM)<sup>c</sup>** |
| --- | --- | --- | --- | --- | --- |
| Wild type | $0.013 \pm 0.002$ | 52 | $0.30 \pm 0.06$ | 2.3 | $\leq 5.8$ |
| Mutant half site | $0.014 \pm 0.001$ | 51 | $1.12 \pm 0.12$ | 0.62 | $\leq 20$ |
| Deleted half site | nd | — | $1.13 \pm 0.20$<sup>d</sup> | 0.61 | — |
| +5 bp insert | nd | — | $0.97 \pm 0.06$ | 0.71 | — |
| +10 bp insert | nd | — | $0.49 \pm 0.01$ | 1.4 | — |

<sup>a</sup> Values are the average of two replicates each using four slide regions, the error is the range.
<sup>b</sup> Half-times were calculated from the average rate constant.
<sup>c</sup> The apparent $K_D$ was calculated using the on and off rate constants and 0.25 nM tetrameric p53.
<sup>d</sup> The average of three replicates, the error is the standard deviation.

**Figure 4.**
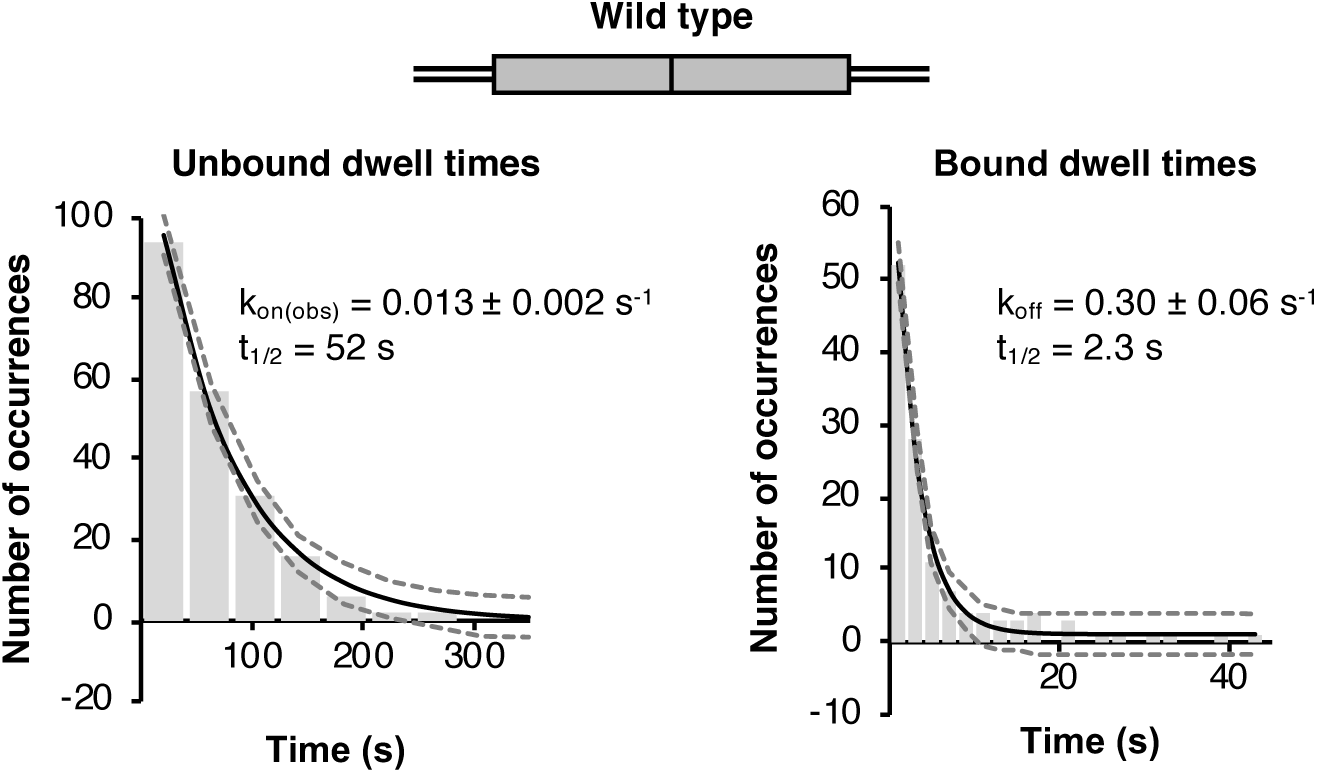
Under equilibrium conditions tetrameric p53 dynamically binds wild type DNA. The unbound (left plot) and bound (right plot) dwell times for p53 tetramer binding events were histogrammed and fit to a single exponential. For the curves shown, the histograms include 208 unbound dwell times and 125 bound dwell times. 95% confidence intervals are represented by the dashed grey lines. The rate constants are the average from two experiments and the errors are the ranges.

We next measured the unbound and bound dwell times for tetrameric p53 on the DNA with a mutant half site. As shown in Figure 5A (left plot), the average k_on(obs)_ was 0.014 ± 0.001 s^-1^. This is not significantly different from the k_on(obs)_ measured on the wild type DNA; hence, mutating one of the half sites did not affect the rate of association (Table 1). On the mutant half site DNA we measured an average k_off_ of 1.12 ± 0.12 s^-1^, corresponding to a half-time of 0.62 sec (Figure 5A, right plot). Therefore, mutating one of the half sites caused tetrameric p53/DNA complexes to be ∼4-fold less stable. This significantly faster off rate on the mutant half site DNA results in lower binding affinity, with an apparent K_D_ of ≤ 20 nM (Table 1). We asked if a similar reduction in the kinetic stability of p53/DNA complexes was observed on the DNA with one half site deleted. Indeed, the average k_off_ on this DNA was 1.13 ± 0.20 s^-1^ (Figure 5B), which is nearly identical to that measured on the mutant half site DNA and ∼4-fold greater than that measured on the wild type DNA. Together our data support the model that p53 tetramers bind different DNAs with the same frequency as long as at least one half site is present; however, the kinetic stability of tetrameric p53/DNA complexes is greater on the DNA containing two half sites (i.e. a full p53 RE) compared to one.

**Figure 5.**
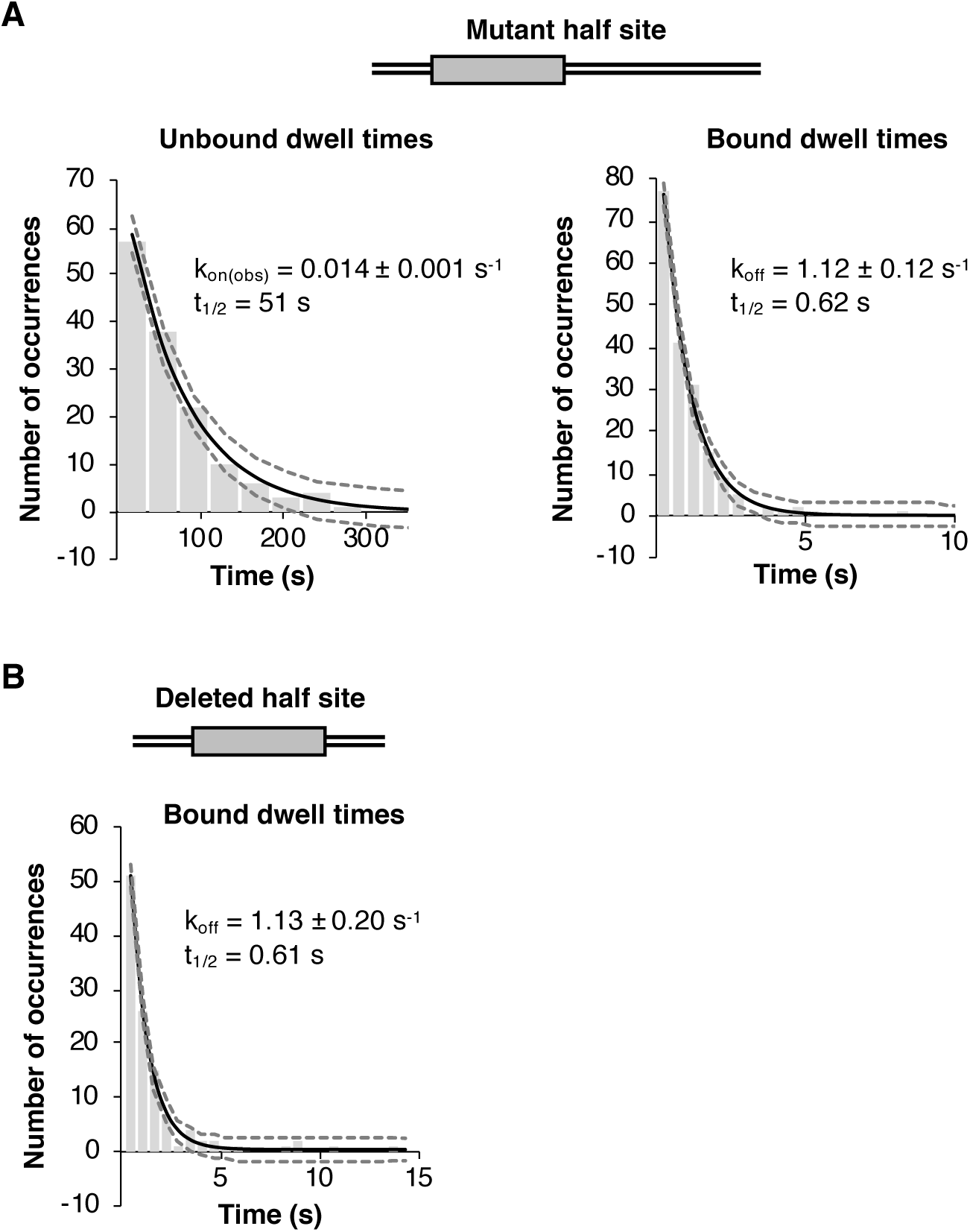
Eliminating one of the DNA half sites reduces the stability of p53/DNA complexes. **A**. Mutating one half site in the DNA does not change the rate of p53 binding, but increases the rate of complex dissociation. Unbound (left plot) and bound (right plot) tetramer dwell times were fit to a single exponential. The histograms shown include 230 unbound dwell times and 186 bound dwell times. The rate constants are the average of two experiments and the errors are the ranges. 95% confidence intervals are represented by the dashed grey lines. **B**. Tetrameric p53 binds to DNA containing a single half site with reduced kinetic stability. Bound tetramer dwell times from three experiments were fit to a single exponential to obtain the average k_off_. The error is the standard deviation. The histogram shown includes 124 bound dwell times.

### Disrupting the helical phasing between half sites destabilizes tetrameric p53/DNA complexes

p53 REs can have variable spacing between the two half-sites, ranging from 0–13 bp (8, 9). Hence, the helical phasing between the two half sites can differ, potentially impacting the binding of p53 to DNA or the kinetic stability of complexes. To test this we designed DNA constructs containing either a 5 bp or 10 bp spacer between the two half sites. As illustrated in Figure 6A, in the wild type DNA with zero spacing between the two half sites, the centers of the two half sites face the same side of the DNA helix. Adding a 5 bp spacer causes the centers of the two half sites to be on opposite faces of the DNA helix, whereas adding a 10 bp spacer restores the helical phasing such that the two half sites are centered on the same face of the DNA helix.

**Figure 6.**
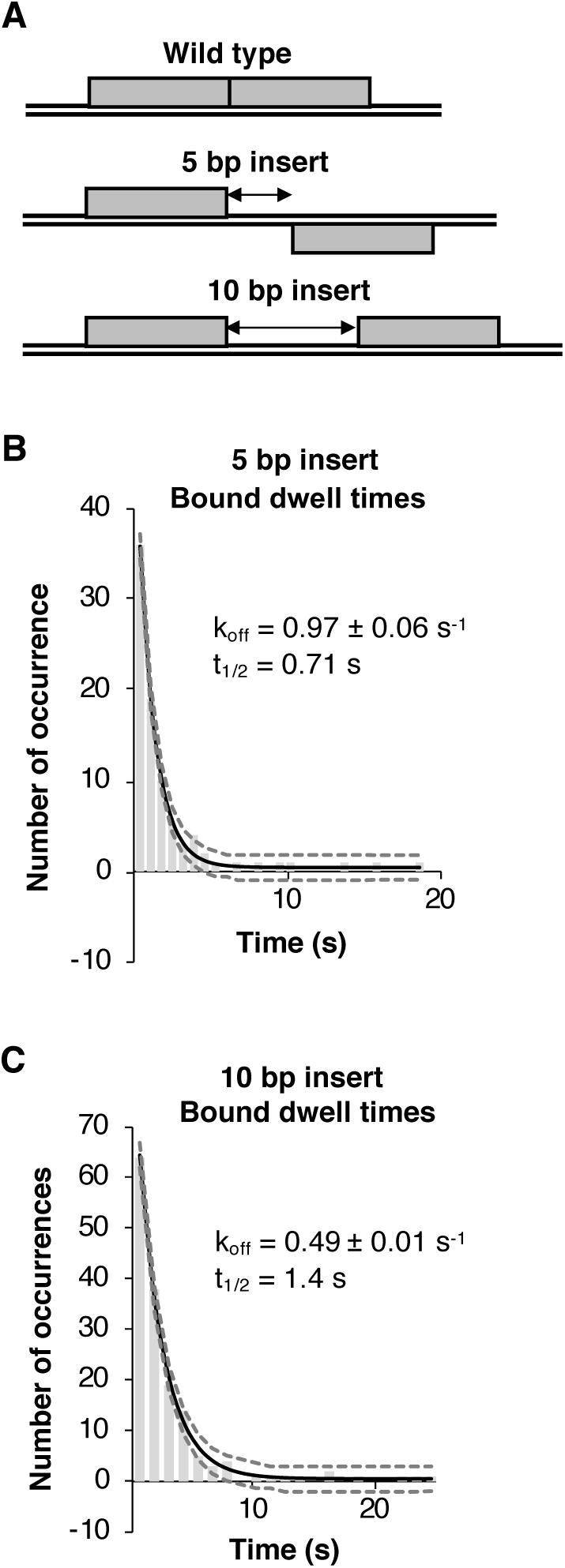
Changing the helical phasing between the half sites impacts the kinetic stability of p53/DNA complexes. **A**. Schematics depicting that in the wild type DNA the half sites are centered on the same face of the DNA helix, insertion of 5 bp between the half sites causes them to be centered on opposite faces of the DNA helix, and insertion of 10 bp results in the half sites being centered on the same face of the DNA helix. **B**. Introducing a 5 bp spacer between the two half sites increases the rate constant for dissociation of tetrameric p53/DNA complexes. The plot shows bound dwell times from 87 states fit to a single exponential. 95% confidence intervals are represented by the dashed grey lines. The k_off_ is the average of two experiments and the error is the range. **C**. The kinetic stability is partially restored when phasing is returned by insertion of 10 bp between the half sites. Shown is a histogram of tetramer bound dwell times from 157 states fit to a single exponential. The k_off_ is the average of two experiments and the error is the range.

We hypothesized that the kinetic stability of tetrameric p53/DNA complexes on the DNA with a 5 bp insert would be similar to that measured on the DNAs with only one half site. We measured the bound dwell times for tetrameric AF647-p53 bound to the 5 bp insert DNA and fit the data to single exponential curve (Figure 6B). We obtained an average k_off_ of 0.97 ± 0.06 s^-1^ (Table 1). This reflects ∼3-fold faster dissociation than we observed with the wild type DNA; moreover, the k_off_ value for the 5 bp insert is not statistically different from those obtained for both the mutant half site and deleted half site DNAs (see Table 1). Hence, centering two half sites on opposite faces of the helix results in p53/DNA complexes that behave similarly to those assembled on DNAs containing only a single half site. We then tested whether restoring the helical phasing between the two half sites would increase the kinetic stability of p53/DNA complexes, using the 10 bp insert DNA. As shown in Figure 6D, we measured an average k_off_ of 0.49 ± 0.01 s^-1^ and a half-time of dissociation of 1.4 sec (Table 1). Hence complexes formed on the 10 bp insert DNA are ∼2-fold more stable than complexes on the 5bp insert DNA. Moreover, the k_off_ on the 10 bp insert DNA is not statistically different from that measured on wild type DNA.

## Discussion

Crucial to the ability of p53 to induce transcription of genes is its recognition and binding to REs. To understand mechanisms by which DNA binding occurs we developed a single molecule fluorescence assay to quantify full length tetrameric p53 binding to DNA in vitro. We found that p53 tetramers bind dynamically and with high affinity to the p53 RE from the GADD45 promoter. When one of the two half sites was removed (via mutation or deletion), tetrameric p53 still bound. With a single half site mutated, the rate of association of tetrameric p53 with the DNA did not significantly change; however, the rate of dissociation of p53/DNA complexes was significantly faster. In addition, when the half sites were centered on opposite faces of the DNA helix, the kinetic stability of tetrameric p53/DNA complexes was similar to that measured with DNAs containing only a single half site. Our real time kinetic measurements of tetrameric p53 binding to different DNAs provide new insight into how this interaction occurs.

Finding that full length p53 binds DNA containing only one half site as a tetramer suggests that one dimer surface binds a half site and the other dimer surface can remain unoccupied. It is possible that p53 tetramers bound to a single half site are structurally different from tetramers bound to two half sites. There are no high resolution structures of full length p53 bound to DNA, likely due to the unstructured N- and C-terminal regions interfering with structure determination. Because these domains impact the ability of p53 to bind DNA (15, 19, 20, 36), it reasonable to predict these regions could impact the structure of p53/DNA complexes. Interestingly, low resolution EM structures with full length p53 suggest two p53 tetramers can simultaneously bind each half site present in a full RE to form an octameric p53 oligomeric state (30–32). We occasionally observed DNAs with higher order p53 oligomers in single molecule experiments (i.e. greater than four dyes bound to DNA); however, the number of these events was a very small fraction of the total binding events and were not considered in our analysis. Hence, we did not obtain strong support for the formation of octameric p53/DNA complexes under the solution conditions in our single molecule experiments.

Our data show that different affinities are measured for p53 binding DNA in EMSAs and in solution single molecule experiments. In EMSAs, p53/DNA complexes formed on wild type and single half site DNAs with apparent sub-nM affinity, and on random DNA with low nM affinity. By comparison, from the single molecule data we calculated K_D_ values for the wild type and mutant half site DNAs of ≤ 5.8 nM and ≤ 20 nM, respectively, and we did not observe any appreciable binding to the random DNA. A likely explanation for the greater apparent affinities in EMSAs is that the native gels trap complexes that are kinetically unstable, leading to an over-estimate of the binding affinity. The K_D_ value we calculated from single molecule measurements with the wild type DNA (GADD45 RE) is comparable to prior K_D_ measurements for this DNA from ensemble fluorescence anisotropy experiments (37).

Our single molecule experiments revealed that p53 binds and releases from DNA dynamically. Indeed, p53 associated with the wild type DNA with a half time of 52 s at the concentration used and had a residence time of 2.3 s. Mutating one of the two half sites did not affect the rate of association of p53 with the DNA, but the stability of the resulting tetramer/DNA complexes was reduced approximately 4 fold, with a residence time of 0.62 s. This loss in kinetic stability is consistent with the model that only one of the dimer interfaces contacts DNA when a single half site is bound by a tetramer. The short residence times of p53 tetramers on the wild type and half site REs are consistent with mounting evidence that transcription factor/DNA associations are dynamic in mammalian cells (38–41). For example, single molecule live cell tracking experiments using fluorescently labeled full length p53 reported average residence times of 3.1–3.5 s (42, 43), which is surprisingly similar to the residence time for p53/wild type DNA that we measured in vitro.

Together our single molecule data provide insight into mechanisms by which p53 tetramers bind DNA. The rates of association of p53 tetramers with wild type and the mutant half site DNAs were indistinguishable, and nearly all binding events occurred in a single step (i.e. without a detectable dimer intermediate). These observations are consistent with the model that p53 tetramers form in solution and then bind DNA. In which case, the rate of association could be set by the rate of formation of p53 tetramers in solution, and that once formed, the tetramers rapidly associate with the DNA. An alternative model is that two dimers bind sequentially to DNA at a rate faster than we can resolve. We believe this model is less likely. Presuming the second binding event is diffusion limited, it would occur over several seconds at the p53 concentration used, which we would readily detect in our assays. In the case of the wild type DNA containing two half sites, the initial recognition and binding by p53 tetramers could be driven by interactions between a single half site and a dimer within tetrameric p53. Then a subsequent conformational change would allow the second dimer to interact with the other nearby half site, causing stabilization of the p53 tetramer/DNA complexes. Conformational transitions are not directly detectable in our single molecule assays; however, our observation that the presence of two half sites significantly stabilized the tetramer/DNA complex is consistent with this model. Moreover, previous studies have suggested a two-step “induced fit” mechanism of binding in which initial DNA binding by p53 is followed by a conformational switch that helps control the dissociation rate (35, 44, 45). Lastly, dissociation of nearly all tetramer p53/DNA complexes occurred without a detectable dimer/DNA intermediate, regardless of DNA construct (Figure 3C). Although it is formally possible that dimers dissociate sequentially at a rate faster than is detectable in our assays.

Our data also revealed a relationship between the kinetic stability of p53/DNA complexes and the spacing between half sites, specifically in regard to their helical phasing. The dissociation of p53 tetramers from DNA containing two half sites on opposite faces of the DNA helix (due to 5 bp spacing) was similar to the dissociation of p53 tetramers from DNA containing only a single half site. This suggests only one of the two half sites was occupied by the p53 tetramer on the DNA with the 5 bp insert. Moreover, a gain in kinetic stability occurred when the half sites were re-centered on the same face of the DNA helix by insertion of 10 bp, suggesting both half sites were contacted by the p53 tetramer. Our kinetic measurements are consistent with a prior report that DNA binding affinity was reduced with 4-5 bp and 14-15 bp spacer lengths, but affinity was largely unaffected with a 10 bp spacer (25). It is possible that a tetramer bound to an RE with 5 bp spacing would have an available binding surface. A prior study found that tetrameric p53 could bind to DNAs with longer spacing between half sites in a “hemispecific” mode, with one p53 dimer bound to a half site and the other dimer bound to the spacer DNA (46).

Although p53 is a widely studied protein, elucidating the mechanism by which it binds DNA is challenging due to the difficulty of studying the full length protein. Here we describe quantitative studies of full length p53 tetramers binding DNA in real time at the single molecule level, providing basic measurements and insight into how this interaction occurs. We are now poised to evaluate how different natural REs and specific regions of p53 contribute to the kinetic mechanisms by which tetramers interact with one and two half sites.

## Experimental Procedures

### Cloning and expression of p53-SNAP protein

We created a fusion construct containing an N-terminal histidine tag, human full length p53, an 8-amino acid linker (GSSGGSSG), and a C-terminal SNAP tag. The p53-SNAP used in the experiment shown in Figure 1A had the amino acid sequence KL prior to the linker. We used PCR with plasmids containing p53 or SNAP sequences and inserted isolated DNA fragments into the pACEBac1 baculovirus transfer vector via EcoRI and XbaI restriction sites. The p53-SNAP pACEBac1 plasmid was transformed into DH10EmBacY cells (Geneva Biotech) and cells were plated on agar plates for blue-white screening (50 µg/mL kanamycin, 10 µg/mL tetracycline, 7 µg/mL gentamycin, 100 µg/mL X-Gal, 40 µg/mL IPTG). A white colony, reflecting the correct transposition of the p53-SNAP insert into the bacmid DNA present within DH10EmBacY cells, was used to inoculate an overnight culture. Bacmid DNA, prone to fragmentation, was prepared from the culture using the following protocol. The cells were pelleted, then resuspended in 330 µL of solution I (15 mM Tris pH 7.9, 10 mM EDTA pH 8, 100 µg/mL RNase A, filter sterilized), followed by addition of 330 µL of solution II (0.2 M NaOH, 1% SDS, filter sterilized). The sample was inverted once to mix and incubated 5 min at room temperature, then 460 µL of 3 M potassium acetate (pH 5.5, autoclaved) was added, gently mixed by inverting, and incubated 5 min on ice. The sample was then centrifuged, the supernatant was transferred to a new tube, and the sample was subjected to a second centrifugation. The supernatant was transferred to a tube containing 900 µL of isopropanol and gently mixed by inverting. After a 15 min incubation on ice, the sample was centrifuged and the supernatant was removed. The pellet was washed with 70% ethanol then air dried for 5 min at room temperature. The pellet was solubilized in TE buffer (10 mM Tris pH 7.9, 1 mM EDTA pH 8, filter sterilized), avoiding mechanical resuspension. The DNA was filtered using a 0.22 µm Spin-X column (Fisher Scientific). Bacmid DNA was sent to the Protein Production/MoAB/Tissue Culture Shared Resource facility at the University of Colorado Anschutz Medical Campus to produce the baculovirus viral stock for infection of SF-9 insect cells to express p53-SNAP. Insect cell pellets were screened for protein expression using western blots against p53, the SNAP tag, and the His tag.

### Purification and fluorescent labeling of p53-SNAP

An SF-9 insect cell pellet expressing full length p53-SNAP was thawed on ice and resuspended in lysis buffer B (50 mM Tris pH 7.5, 150 mM NaCl, 10% glycerol, 0.5% NP-40, 2 mM DTT, 1 mM PMSF, 1 µg/mL Pepstatin A, 1X EDTA-free Protease Inhibitor (Roche), 1X PhosSTOP (Roche)) by vortexing then incubating on ice for 15 min before centrifugation. The supernatant was loaded onto a HisPur Ni-NTA resin (Thermo Scientific) column that was pre-equilibrated with lysis buffer B. The column was then washed sequentially with lysis buffer B, wash buffer A (50 mM Tris pH 7.5, 500 mM NaCl, 5 mM MgCl_2_, 10% glycerol, 2 mM DTT, 1 mM PMSF, 1 µg/mL Pepstatin A, 1X EDTA-free Protease Inhibitor cocktail (Roche), 1X PhosSTOP (Roche)), and wash buffer A containing 50 mM imidazole. p53-SNAP was eluted with wash buffer A containing 300 mM imidazole. Fractions were analyzed via SDS-PAGE, and the fractions containing the most p53-SNAP were pooled and dialyzed overnight in 50 mM Tris pH 7.9, 150 mM NaCl, 5 mM MgCl_2_, 10% glycerol, and 1 mM DTT.

For the protein labeling reaction, 100 µL of p53-SNAP (diluted to a concentration less than 10 µM) was incubated with 2 µL of 1 mM SNAP-Surface^®^ Alexa Fluor^®^ 647 dye (New England Biolabs) for 2 hr at room temperature in the dark. Excess free dye was removed using Zeba Spin Desalting columns (7K MWCO, 0.5 mL, Thermo Scientific). A coomassie stained SDS-PAGE gel with a BSA standard curve was used to calculate the total concentration of p53-SNAP. The concentration of AF647-p53 was evaluated by comparison to a standard curve of AF647-SNAP substrate and a standard curve of an AF647-labeled oligonucleotide using a Typhoon 9500 Imager. The absorbance at 652 nm and the molar absorptivity of AF647 was also used to calculate the concentration of AF647 in the labeled p53. In addition, LC-MS/MS was used to measure the amount of the 16-amino acid peptide (TALSGNPVPILIPCHR) containing the site of AF647 dye attachment in trypsin digested samples of unlabeled p53-SNAP and AF647-p53. The peptide was not detectable in the AF647-p53 sample, but readily detectable in the p53-SNAP sample, indicating complete conjugation to the dye.

### Oligonucleotides

The sequences of the DNA constructs used in binding assays were as follows (ordered HPLC purified from Integrated DNA Technologies): Wild type reverse oligo, 5’CGACGCCAGCATGCTTAGACATGTTCGCTCTA3’ and Wild type forward oligo, 5’TAGAGCGAACATGTCTAAGCATGCTGGCGTCG3’; Mutant half site reverse oligo, 5’GACGCCGCCTTTGAAAGACATGTTCGCTCTA3’ and Mutant half site forward oligo, 5’TAGAGCGAACATGTCTTTCAAAGGCGGCGTCG3’; Deleted half site reverse oligo, 5’CGACGCAGACATGTTCGCTCTA3’ and Deleted half site forward oligo, 5’TAGAGCGAACATGTCTGCGTCG3’; Random DNA reverse oligo, 5’TATTCGTGTCAGCCCCTACCCGATTAGGAACG3’ and Random DNA forward oligo, 5’CGTTCCTAATCGGGTAGGGGCTGACACGAATA3’; 5bp insert reverse oligo, 5’CGACGCCAGCATGCTTTCCACAGACATGTTCGCTCTA3’ and 5bp insert forward oligo, 5’TAGAGCGAACATGTCTGTGGAAAGCATGCTGGCGTCG3’; 10 bp insert reverse oligo 5’CGACGCCAGCATGCTTATTAATTCATAGACATGTTCGCTCTA3’ and 10 bp insert forward oligo 5’TAGAGCGAACATGTCTATGAATTAATAAGCATGCTGGCGTCG3’. For single molecule experiments, all reverse oligos contained a AF647 dye on the 5’ end and all forward oligos contained a 5’ biotin attached to a 24 nt single stranded linker: 5’CGCGTTCATGGTAGAGTCGTGGAC3’. For EMSAs, all reverse oligos were radiolabeled on the 5’end using T4 polynucleotide kinase and γ^32^P^-^ATP.

Double stranded DNAs were generated by annealing the forward and reverse oligos by heating to 95° C for 10 min and slowly cooling to room temperature in the temp block on the bench top. For single molecule experiments, annealed DNAs were gel purified on a native 7% polyacrylamide gel (containing 0.5X TBE, pre-run at 150V for 30 min prior to loading DNA samples). DNAs were cut out of the gel, slices were crushed in TE-Low buffer (10 mM Tris pH 7.9, 0.1 mM EDTA), then nutated overnight nutation at 4° C. DNA was isolated using Spin-X columns (Fisher Scientific) and ethanol precipitation.

### Electrophoretic mobility shift assays (EMSAs)

DNA binding reactions (20 µL) were assembled with wild type p53, p53-SNAP, or AF647-p53 (concentrations noted in figure legends) and ^32^P-DNA (0.1 nM for Figure 1A and 0.03 nM for Figures 1C and 3B) in 25 mM HEPES pH 7.9, 50 mM KCl, 2.5 mM MgCl_2_, 1 mM EDTA, 10% glycerol, 0.1% NP-40, 1 mM DTT and 0.05 mg/mL BSA. Reactions were incubated for 30 min at room temperature. Samples were run on 4% native polyacrylamide gels (containing 0.5X TBE, pre-run at 150V for 30 min) in 0.5X TBE running buffer at 150V for 1 hr. Gels were then dried and imaged on a Typhoon 9500 Imager.

### Single molecule binding assays and data collection using TIRF microscopry

Cleaning and assembly of the flow chambers and preparation of stock solutions of streptavidin, D-glucose, glucose oxidase, catalase, and 100 mM Trolox (Sigma) for single molecule microscopy were as previously described (47). Flow chambers were washed twice with MilliQ water and 1X buffer (25 mM Tris 7.9, 50 mM KCl, 5 mM MgCl_2_. 10% glycerol, 0.05 mg/mL BSA, 1 mM DTT, and 0.1% NP-40) prior to use. The surface was then incubated with 0.2 mg/mL streptavidin (Sigma) and 0.8 mg/mL BSA diluted in 1X buffer for 5 min at room temperature. Excess streptavidin was washed out with additional 1X buffer. 10 pM of biotinylated DNA was then flowed onto the surface and incubated for 10 min to allow immobilization. Excess DNA was washed out with additional 1X buffer. Imaging buffer (1.02 mg/mL glucose oxidase, 0.04 mg/mL catalase, and 0.83% D-glucose diluted in 1X buffer containing 3.45 mM Trolox) was then flowed into the chamber. DNA emission movies were collected over four regions with the use of a piezo nanopositioning stage. AF647-p53 (0.25 nM tetramer) in imaging buffer was flowed into the chamber and incubated for 10 min. Without washing, p53+DNA emission movies were collected over the same regions as the DNA emission movies using the piezo stage.

Emission movies were collected with a 1.49 NA immersion objective based TIRF microscope (Nikon TW-2000U) equipped with two CCD cameras. Emission from the 635 nm laser with a power setting of 108 mW was recorded by a Cascade II Photometric CCD camera using the NIS-elements software. Emission movies of DNA only were collected with a 60 ms exposure time for 100 frames. Emission movies of p53+DNA were collected with a 60 ms exposure for 1000 frames to obtain bound dwell times. To obtain unbound dwell times on wild type DNA the data were collected with a 200 ms exposure for 2000 frames, and a 600 ms exposure for 1000 frames; for the mutant half site DNA these parameters were 60 ms for 5000 frames.

### Analysis of single molecule data

In-house software was used to co-localize spots in the DNA and p53+DNA movies, quantify the oligomeric state of bound p53, and extract dwell times. Briefly, for a spot pair found between the DNA movie and p53+DNA emission movie, the intensity trace from the DNA emission movie was used to calculate a base red intensity N that corresponded to the value of a single AF647 dye. The value N was used to calculate the number of dyes present for each state in the p53+DNA intensity trace of the spot pair. The total number of dyes was used to characterize a state as being unbound (one dye) or bound by tetrameric p53 (four or five total dyes). Transitions between different intensity states representing dissociation or association of p53 tetramers were identified. The number of unbound DNA states, dynamic tetrameric p53-DNA bound states, bound dwell times, and unbound dwell times between states were calculated then compiled for all regions for a given experiment. Spots showing bound p53 with no transitions across the time of collection were not included in subsequent analyses.

## Acknowledgments

This research was funded by grant MCB-1817442 from the National Science Foundation. E.L. was also supported by NSF Graduate Research Fellowship Program award DGE1144083 and training grant T32 GM08759 from the National Institutes of Health.

## Conflict of Interest

The authors declare that they have no conflicts of interest with the contents of this article.

## Abbreviations

RE: p53 response element;
EMSA: electrophoretic mobility shift assay;
DBD: DNA binding domain;
OD: oligomerization domain;
CTD: C-terminal domain, p53-SNAP, p53 containing a SNAP tag;
AF647-p53: p53-SNAP labeled with AlexaFluor647;
TIRF: total internal reflection fluorescence microscopy.

